# The absence of SOX2 in the anterior foregut alters the esophagus into trachea and bronchi in both epithelial and mesenchymal components

**DOI:** 10.1101/739714

**Authors:** Machiko Teramoto, Ryo Sugawara, Katsura Minegishi, Masanori Uchikawa, Tatsuya Takemoto, Atsushi Kuroiwa, Yasuo Ishii, Hisato Kondoh

## Abstract

In the anterior foregut (AFG) of mouse embryos, the transcription factor SOX2 is expressed in the epithelia of the esophagus and proximal branches of respiratory organs comprising the trachea and bronchi, whereas NKX2.1 is expressed only in the epithelia of respiratory organs. Previous studies using hypomorphic *Sox2* alleles have indicated that reduced SOX2 expression causes the esophageal epithelium to display some respiratory organ characteristics. In the present study, we produced mouse embryos with AFG-specific SOX2 deficiency. In the absence of SOX2 expression, a single NKX2.1-expressing epithelial tube connected the pharynx and the stomach, and a pair of bronchi developed in the middle of the tube. Expression patterns of NKX2.1 and SOX9 revealed that the anterior and posterior halves of SOX2-deficient AFG epithelial tubes assumed the characteristics of the trachea and bronchus, respectively. In addition, we found that mesenchymal tissues surrounding the SOX2-deficient NKX2.1-expressing epithelial tube changed to those surrounding the trachea and bronchi in the anterior and posterior halves, as indicated by the arrangement of smooth muscle cells and SOX9-expressing cells and by the expression of *Wnt4* (esophagus specific), *Tbx4* (respiratory organ specific), and *Hoxb6* (distal bronchus specific). The impact of mesenchyme-derived signaling on the early stage of AFG epithelial specification has been indicated. Our study demonstrated an opposite trend where epithelial tissue specification causes concordant changes in mesenchymal tissues, indicating a reciprocity of epithelial-mesenchymal interactions.

## Introduction

Epithelial components of the vertebrate alimentary tract develop from the endoderm-derived gut tube, which becomes specified into subdivisions along the anteroposterior axis. Early-stage subdivisions of endoderm-derived gut tubes depend on the action of various exogenous signaling molecules (Jacobs et al., 2012; Swarr and Morrisey, 2015; Zhang et al., 2017), and specification to subdivision results in the expression of subdivision-specific transcription factors (TFs). The epithelial tube of the anterior foregut (AFG) and stomach expresses SOX2 after E9 (Que et al., 2007), whereas the hindgut, which develops into duodenum and more posterior intestinal regions, expresses CDX2 (Gao et al., 2009). The anterior portion of AFG epithelial tubes is then split into dorsally positioned SOX2-expressing esophagus tissues and ventrally positioned tracheal tissues that express NKX2.1 and SOX2. Bronchi, bronchioles, and alveoli that further develop at the distal end of the trachea also express NKX2.1 (Swarr and Morrisey, 2015; Zhang et al., 2017).

TFs that are expressed in endoderm-derived epithelial tubes play determining roles during gut subdivision. For instance, endoderm-specific inactivation of *Cdx2* promoted SOX2 expression in hindgut tubes, which assumed forestomach epithelial cell characters (Gao et al., 2009). Loss of *Nkx2*.*1* in embryos resulted in the development of esophagus and tracheal structures that remained joined, and caused defects in lung branching morphogenesis (Minoo et al., 1999).

Contributions of SOX2 to esophageal specification of AFG epithelia have been demonstrated using combinations of hypomorphic *Sox2* alleles, leading to degrees of tracheoesophageal fistula (Que et al., 2007), similar to various aspects of human congenital tracheoesophageal fistula cases (Wong NC, 2016; Zhang et al., 2017). Hence, SOX2 function is essential for the separation of trachea and esophagus in the AFG and required for the establishment of esophageal epithelial characteristics. However, complete tissue-specific *Sox2* inactivation is required to precisely evaluate the roles of SOX2 in these processes. We achieved endoderm-specific inactivation of *Sox2* using a floxed *Sox2* allele and a *FoxA2*-*nEGFP*-*CreERT2* knock-in allele, in which coding sequences were joined via 2A peptide sequences (Imuta et al., 2013). Under these conditions, tamoxifen administration resulted in the development of a single SOX2-deficient AFG tube connecting the pharynx and the stomach, in the middle of which a pair of bronchi branched out.

Mesenchymal influences on epithelial subdivisions of the AFG have been well documented (Swarr and Morrisey, 2015; Zhang et al., 2017). In this study, we characterized both epithelial and mesenchymal components of the AFG after formation in the absence of SOX2 in the epithelium. Not only the SOX2-deficient AFG epithelia but also the surrounding mesenchymal tissues developed into those of respiratory organs, with clear anteroposterior polarity similar to those of the trachea and bronchi. Thus, once established, the regional identity of the epithelial component of the AFG determined the characters of the surrounding mesenchymal tissues.

## Results and Discussion

### Organization of AFG tube development in the absence of endodermal SOX2 expression

To inactivate *Sox2* expression in the entire gut tube, we introduced *Foxa2*^*nEGFP-CreERT2*^ (Imuta et al., 2013) into floxed *Sox2* homozygous mice (Fig. S1A–C). Tamoxifen was administered to pregnant females on embryonic days 7 and 8 (E7 and E8), and embryos were collected on E11 to E13. In the floor plate-proximal ventral spinal cord, where the expression of *Sox2* and *FoxA2* normally overlaps, SOX2 expression was abolished (Fig. S1D), confirming that *Sox2* was efficiently inactivated by CreERT2.

We investigated AFG development in the absence of SOX2 expression (Fig. 1). AFG epithelial tubes were visualized using *FoxA2*-driven EGFP fluorescence (Fig. 1A; Fig. S2). In normal E11 to E13.5 embryos, the esophagus joins the pharynx and stomach, and the trachea with a pair of bronchi with lung tissues was already separated (Fig. 1A). In the absence of SOX2 expression, however, a single AFG tube connecting the pharynx and stomach was formed, and a pair of bronchi with lung tissues developed in the middle of these tubes (Fig. 1A). In a significant fraction (12/17) of SOX2-deficient AFG, bronchiole-like tissues developed from the posterior half (Fig. 1A; Fig. S2, arrows). The same AFG tube phenotype was observed using knock-in of CreERT2 into the ROSA26 locus (Cheng et al., 2010), although in this case, brain tissue development was also affected (data not shown). Loss of SOX2 expression in endoderm-derived epithelia was confirmed by immunostaining of cross sections (Fig. 1B).

**Figure 1.**
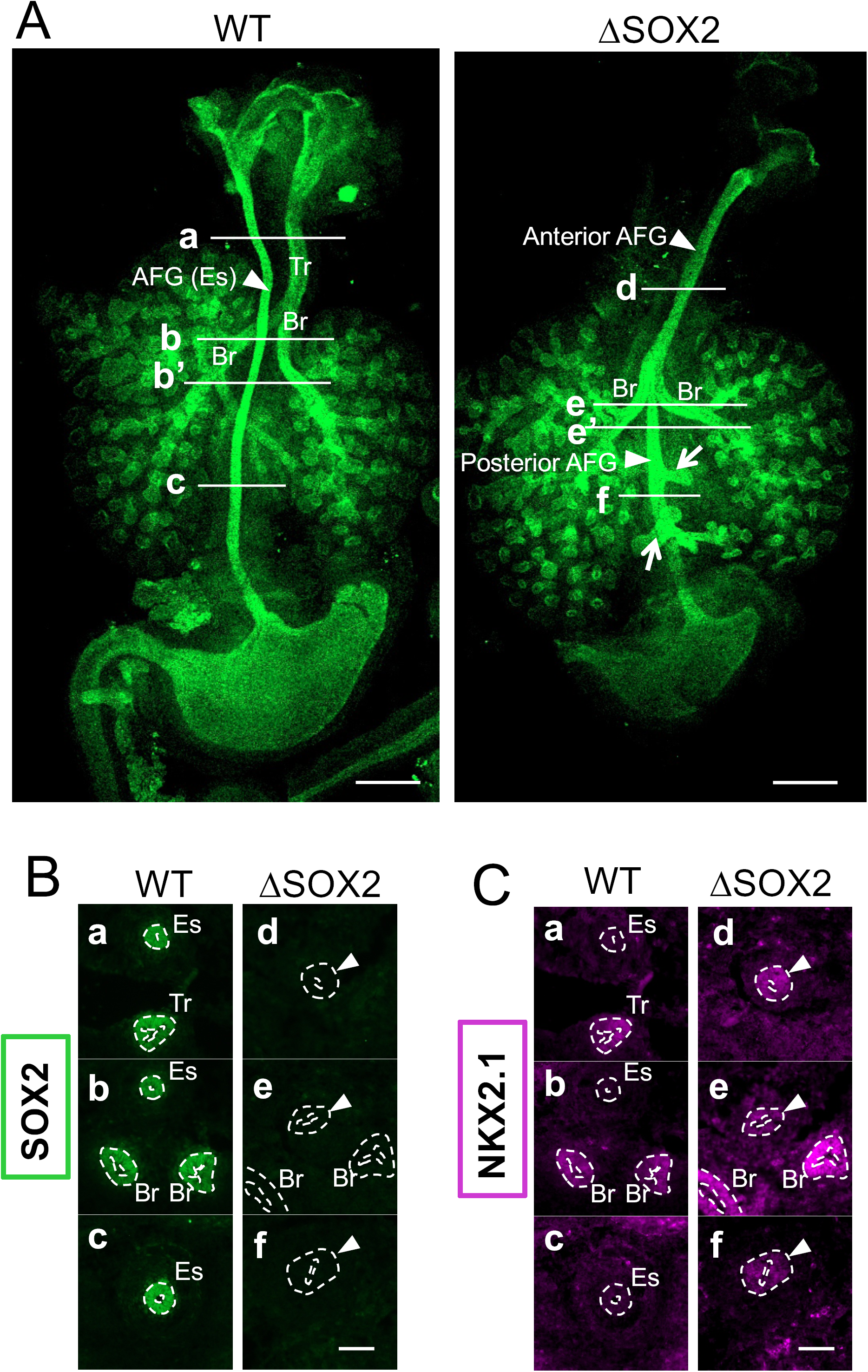
Absence of the esophagus and expression of NKX2.1 in all epithelial tubes of SOX2-deficient (ΔSOX2) AFG. **(A)** AFG tube organization was visualized in the dorsal view by *FoxA2*-driven EGFP in WT AFG and ΔSOX2 AFG on E13.5. WT AFG had separate esophageal (Es) and tracheal (Tr)/bronchial (Br) tubes, whereas ΔSOX2 AFG bronchi developed in the middle of the single tube connecting the pharynx and stomach. Frequently (12/17), bronchiole-like tissues developed from the posterior half of ΔSOX2 AFG, as indicated by arrows. Stomachs (St) of ΔSOX2 were always smaller. Bars, 500 µm. For more specimens from earlier developmental stages, see Fig. S2. **(B and C)** Immunostaining for SOX2 and NKX2.1, respectively, of cross sections of AFG tubes on E13 made at approximate axial levels as shown in **(A)**. Arrowheads indicate the ΔSOX2 AFG epithelium, while Es, Tr, and Br indicate esophagus, trachea, and bronchi, respectively. In ΔSOX2 AFG, the entire epithelial tube lacked SOX2 and expressed NKX2.1. See also Fig. S3. Bars, 100 µm.

In normal embryos, proximal branches of the epithelial tubes, trachea, bronchi, and bronchioles, also express SOX2 (Fig. 1B; Fig. S3A) (Gontan et al., 2008; Que et al., 2009; Wong NC, 2016). Airway epithelia that developed in the absence of SOX2 showed tissue growth and branching patterns identical to normal embryos (Fig. 1A; Fig. S2), indicating that SOX2 function is dispensable for lung morphogenesis at least up to the E13 stage. It has been shown that SOX2 is involved in later stages of lung development (Que et al., 2009).

SOX2-deficient stomach tissues were smaller than normal (n>10) (Fig. 1A; Fig. S2), but did not express NKX2.1 (n=3) (Fig. S3B). CDX2 is normally expressed in hindgut epithelia to form SOX2–CDX2 boundaries at the pylorus. In the absence of CDX2 expression, hindgut epithelia expressed SOX2 and assumed a forestomach character (Gao et al., 2009). In contrast, the absence of SOX2 expression did not activate CDX2 expression in the stomach epithelia (n=3) (Fig. S3C).

### SOX2-deficient single AFG epithelial tubes have tracheal and bronchial characters

SOX2-deficient AFG epithelial tubes connecting the pharynx and stomach expressed NKX2.1 at all axial levels during the stages E11 to E13.5 (n=4) (Fig. 1C and Fig. S3AB), indicating tracheal/bronchial characteristics. Accordingly, on E13.5, SOX2-deficient AFG tubes comprising columnar (Co) epithelia were wider than normal esophagi (Fig. 2A), again showing tracheal and bronchial characteristics, whereas wild-type (WT) esophagi comprised pseudo-stratified (PS) epithelial tissues at this stage (Fig. 2A), a transition state between the earlier columnar and later stratified epithelia of the esophagus (Que, 2015).

**Figure 2.**
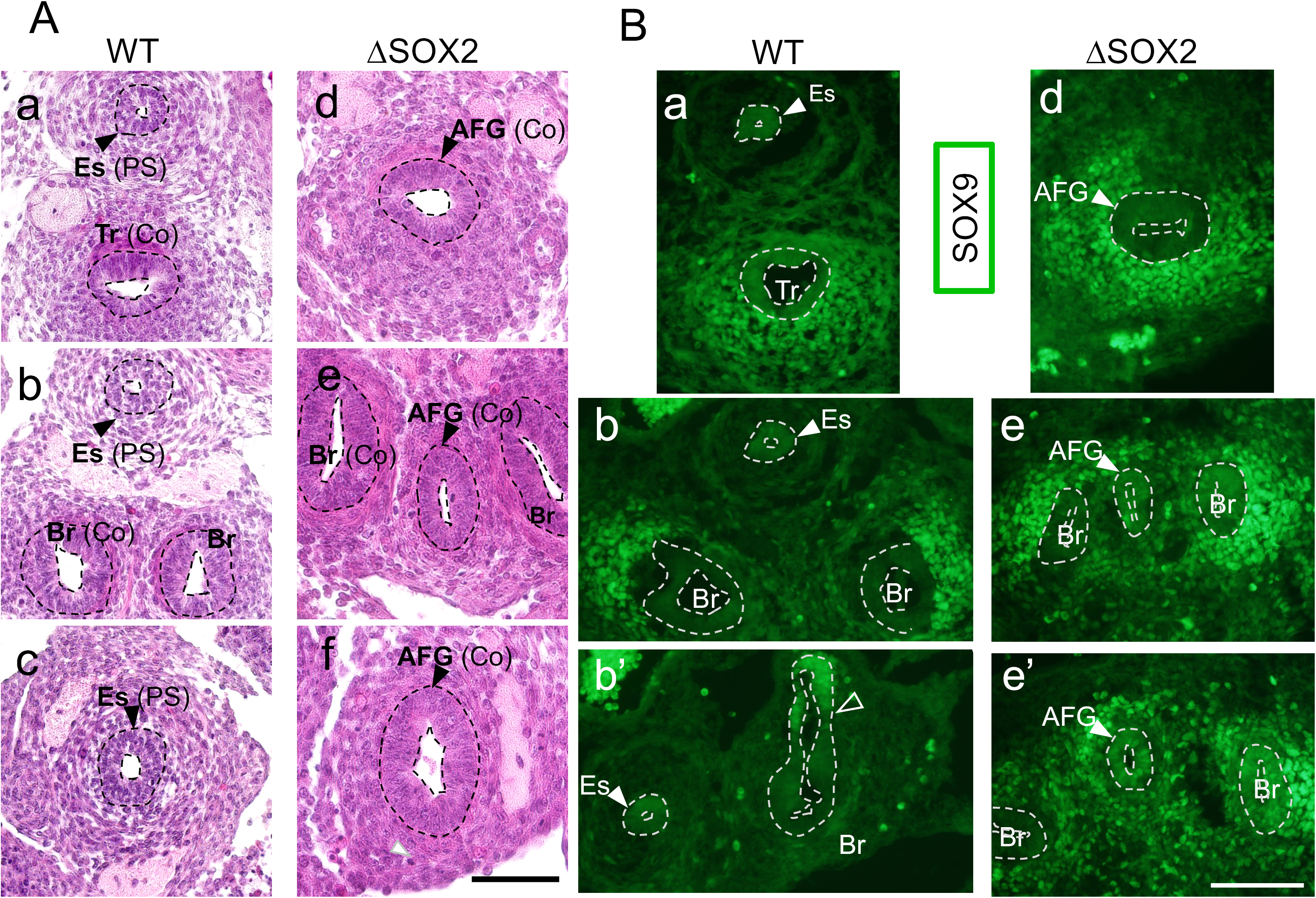
Epithelial tubes in ΔSOX2 AFG exhibit tracheal/bronchial characters. Histological sections of E13 epithelial tubes were made at the axial levels shown in Fig. 1A. **(A)** H&E staining of embryo cross sections. WT esophageal tubes (Es) comprised pseudo-stratified epithelium (PS), whereas WT tracheal (Tr)/bronchial (Br) tubes and ΔSOX2 AFG tubes comprised columnar (Co) epithelium. **(B)** Immunostaining of cryosections for SOX9 at approximate axial levels shown in Fig. 1A. SOX9 expression level was high in posterior ΔSOX2 AFG tubes (e*’*), similar to the distal bronchi in both WT and ΔSOX2 embryos. The open arrowhead indicates the region of transition from low to high SOX9 expression. Bars, 100 µm.

On E13, proximal and distal epithelial portions of respiratory organ rudiments differ in SOX9 expression levels (Wong et al., 2016): low in the trachea and proximal bronchi (Fig. 2Ba,b) and high in the distal bronchi (Fig. 2Bb’). In contrast, in WT esophagi, SOX9 expression did not differ significantly between the anterior and posterior halves (n=5), although the expression levels were variable between embryos and sometimes only barely detectable. In SOX2-deficient AFG tubes, epithelial SOX9 expression was low in the anterior part (Fig. 2Bd,e) but higher in a posterior part (n=4) (Fig. 2Be*’*), indicating that the anterior and posterior sides of the AFG developed as trachea and bronchi, respectively. Given that ordinary pairs of bronchi branched out from the midpoint of SOX2-deficient AFG tubes, the AFG region between the midpoint and the stomach developed as a third bronchus. Consistently, additional bronchiole-like tissues frequently developed from the posterior half of SOX2-deficient AFG (n=12/17) (Fig. 1A; Fig. S2, arrows).

### Surrounding mesenchymal tissues developed concordant with the epithelial tissue identity

We investigated whether mesenchymal tissues surrounding SOX2-deficient AFG tubes developed concordantly with the tracheal and bronchial characteristics of the epithelia.

In normal embryos on E13, the epithelial tubes of the esophagus were surrounded by smooth muscle cells expressing smooth muscle α-actin (α-SM) at positions distal from the tube (Fig. 3A) (McHugh, 1995) and were not associated with SOX9-expressing cells (Fig. 3A). In contrast, the epithelial tubes of the trachea and bronchi were tightly wrapped in two patches of mesenchymal cells, representing smooth muscle cells and SOX9-expressing chondrocyte progenitors. These smooth muscle patches were positioned at the dorsal face of the trachea and median faces of the bronchi and were broader surrounding the bronchi (Fig. 3A). The anterior halves of SOX2-deficient AFG tubes were also tightly wrapped in a narrower patch of smooth muscle cells and a broader patch of SOX9-expressing cells, exactly as observed in normal trachea (n=3) (Fig. 3Aa,d). These data indicate that trachea-like development of the anterior epithelial tube elicits mesenchymal developments with characteristics of respiratory organs.

**Figure 3.**
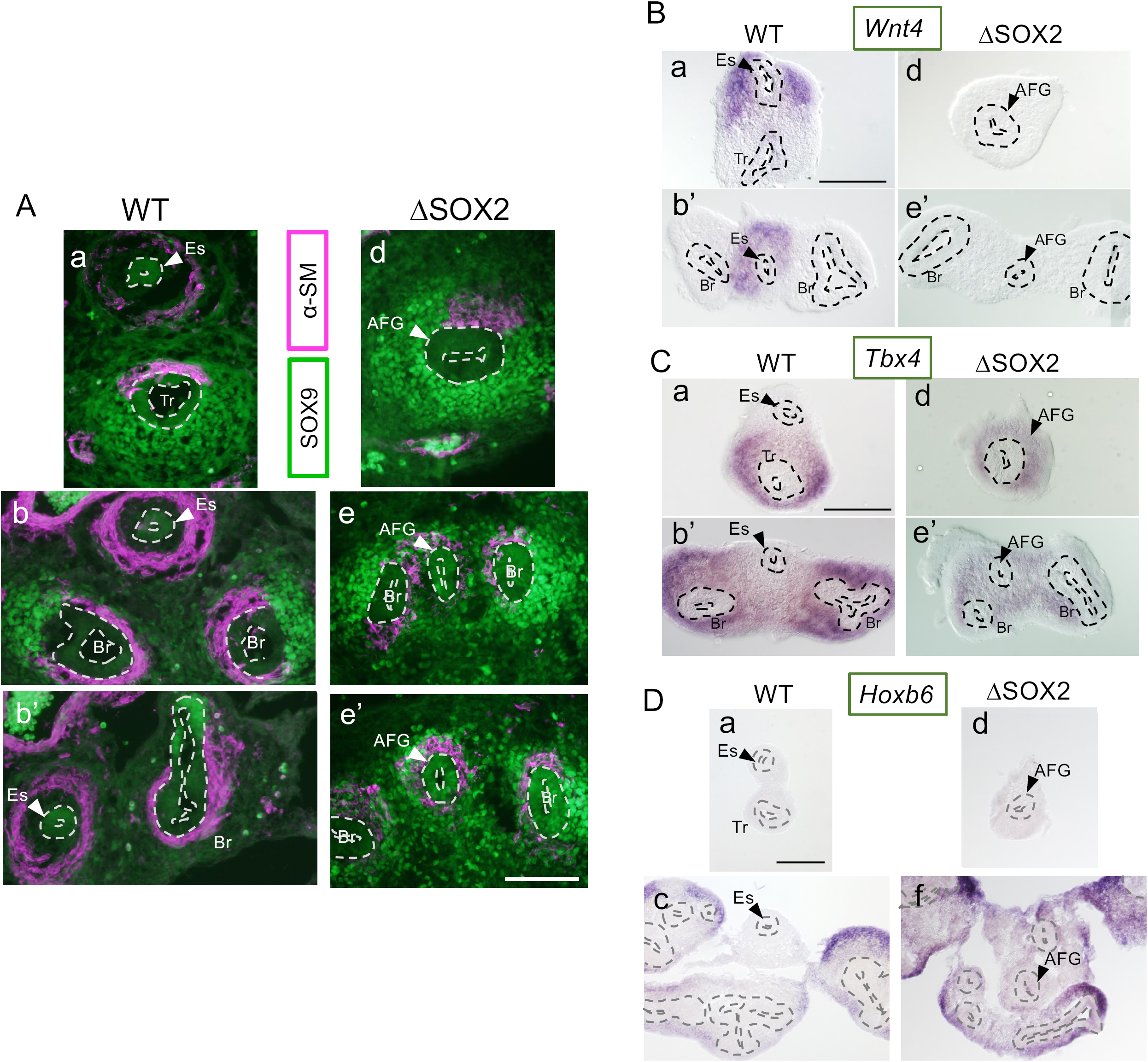
Mesenchymal characters are concordant with those of epithelial tubes in the AFG. Histological sections of E13 epithelial tubes are shown at the axial levels indicated in Fig. 1A. **(A)** Sections shown in Fig. 2B were doubly stained for SOX9 (green) and α-SM (magenta). Mesenchymal tissues surrounding WT and ΔSOX2 trachea/bronchi comprise nearly complementary patches of SOX9- and α-SM-expressing cells. ΔSOX2 AFG tubes were also wrapped in SOX9- and α-SM-expressing mesenchymal tissues, albeit without their separation into patches. In contrast, WT esophageal tissues lacked these cell types and only had distal α-SM-expressing cells. **(B–D)** Analogous sections of WT and ΔSOX2 AFG hybridized with mesenchyme-specific gene probes for *Wnt4* (esophagus specific), *Tbx4* (trachea/bronchus specific), and *Hoxb6* (distal bronchus specific). In ΔSOX2 AFG, *Wnt4* expression was completely absent **(B)**, epithelial tubes were all surrounded by *Tbx4*-expressing mesenchyme **(C)**, and the posterior part of the tube was surrounded by *Hoxb6-*expressing mesenchyme, similar to the distal bronchi **(D)**. Bars, 100 µm.

In the posterior parts of SOX2-deficient AFG, the epithelial tubes were surrounded by smooth muscle cells and SOX9-expressing cells, resembling bronchi. Yet, smooth muscle cells and SOX9-expressing cells did not segregate (n=3) (Fig. 3Ae,e*’*). Presumably, azimuthal specification of the mesenchyme failed to occur in SOX2-deficient AFG.

To investigate the molecular markers of mesenchymal cells that distinguish esophagus and respiratory organs, we performed *in situ* hybridization using esophagus mesenchyme-specific *Wnt4* (Fig. 3Ba,b*’*) (GSE118641, Gene Expression Omnibus), respiratory organ-specific *Tbx4* (Fig. 3Ca,b*’*) (Arora et al., 2012), and *Hoxb6*, which is expressed in the distal portion of the bronchus mesenchyme (Fig. 3Dd).

In normal embryos on E11 to 11.5, *Wnt4* was expressed in mesenchymal cells surrounding the esophagus (Fig. 3Ba,b*’*). In contrast, *Wnt4* expression was lost in mesenchymal tissues surrounding SOX2-deficient AFG epithelial tubes (n=3) (Fig. 3Bd,e*’*), confirming the loss of esophageal phenotypes. In contrast, *Tbx4* was expressed exclusively in the mesenchyme of respiratory organs in normal embryos (Fig. 3Ca,b*’*) (Arora et al., 2012). Mesenchymal tissues of SOX2-deficient AFG expressed *Tbx4* at all axial levels (n=3) (Fig. 3Cd,e*’*), concordant with the respiratory organ identity of the epithelial tubes.

In normal embryos, *Hoxb6* was expressed in mesenchymal tissues surrounding the distal part of the bronchus (and more strongly in alveoli) (Fig.3Dc), but no expression was observed along the trachea (Fig. 3Da). In SOX2-deficient AFG tubes, however, only the posterior region of the mesenchyme expressed *Hoxb6* (n=4) (Fig. 3Df), as observed in the distal bronchi (Fig. 3Dc,f). These observations confirmed the bronchial identity of the posterior half of SOX2-deficient AFG, complete with proximodistal polarity in both epithelial and mesenchymal components (Figs. 2B and 3D).

### Overall impact of the absence of SOX2 during esophageal and tracheal/bronchial development

We investigated the consequences of abolishing SOX2 expression in the anterior endoderm. Under this condition, single AFG epithelial tubes connecting the pharynx and stomach developed, comprising columnar epithelial cells and expressing NKX2.1, indicating gain of tracheal/bronchial characteristics and loss of esophageal characteristics (Figs. 1 and 4). Trisno et al. (2018) similarly ablated *Sox2* in the endoderm and reported the development of NKX2.1-expressing AFG epithelia that lacked esophagus-characteristic p63 expression.

**Figure 4.**
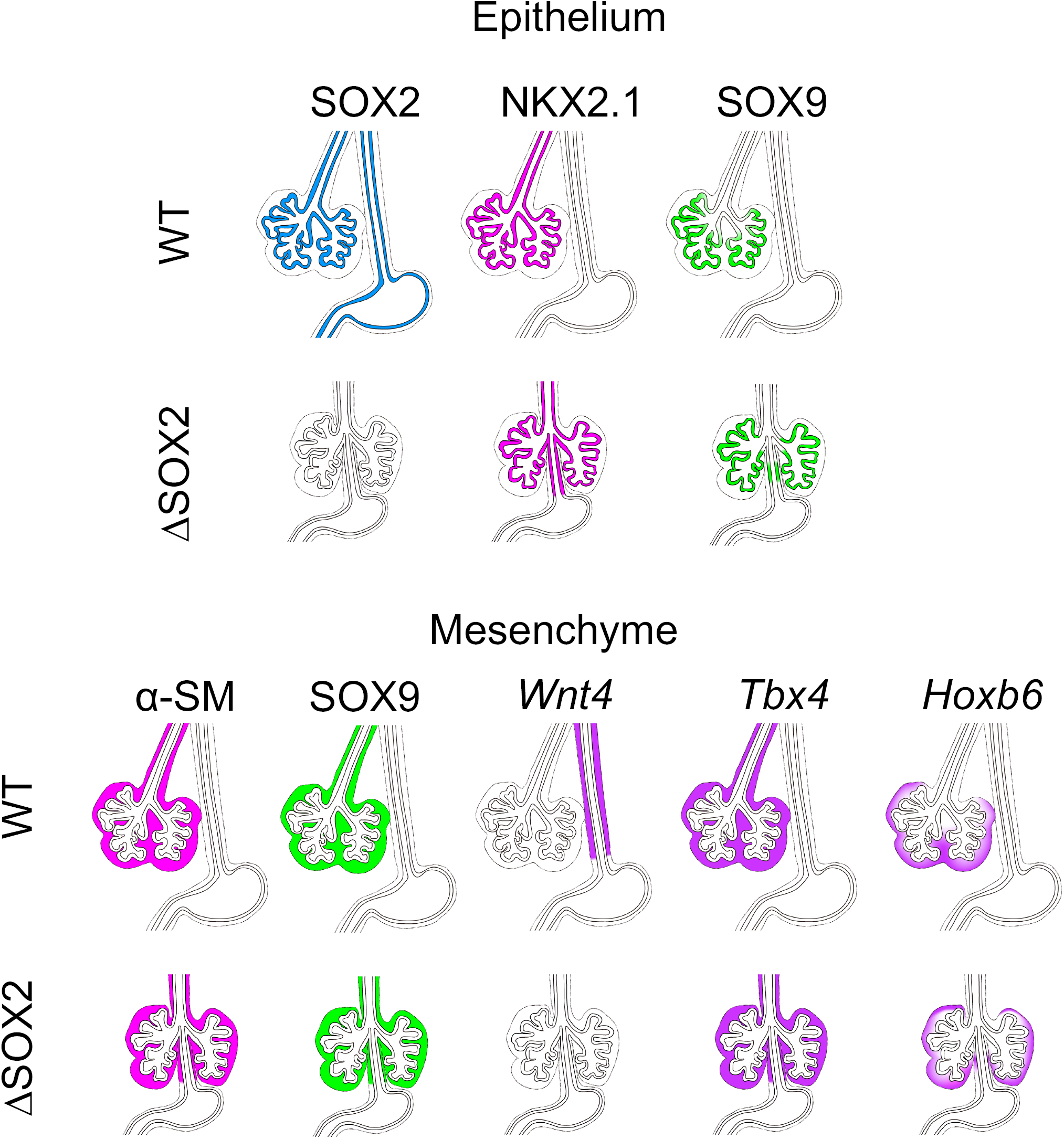
Summary of the key observations of this study. In the diagrams of WT and ΔSOX2 AFG tubes, expression of SOX2, NKX2.1, and SOX9 in the epithelial tissues and expression of α-SM, SOX9, *Wnt4, Tbx5*, and *Hoxb6* in mesenchymal tissues are indicated. These data indicate that the loss of SOX2 in the epithelial tissues alters the esophageal characters of the AFG into trachea and bronchi in both epithelial and mesenchymal components.

We also observed anteroposterior differentiation of SOX2-deficient AFG epithelial tubes, similar to proximodistal differentiation of respiratory organ epithelia (Figs. 2B). The anterior part of SOX2-deficient AFG epithelial tubes had low SOX9 expression, as in trachea, whereas a posterior part of the tube had high SOX9 expression, as observed in the distal bronchi.

A new important finding of this study is that mesenchymal tissues that surround the epithelial tubes change their characters at all axial levels in concordance with the changes in the epithelial tubes from the esophagus to the respiratory organs (Figs. 3 and 4). Studies of alimentary tract development have provided a paradigm of epithelial-mesenchymal interactions during organ development (Swarr and Morrisey, 2015; Zhang et al., 2017). Earlier studies demonstrated the impact of mesenchyme-derived cues on the development of epithelial tubes. For example, dorsoventral differentiation during the early stages of AFG development, leading to esophagus/trachea splitting, is mediated by BMP signaling from the surrounding mesenchymal cells (Domyan et al., 2011; Ishii et al., 1998). Yet, regulatory influences in SOX2-deficient AFG epithelial tubes were in the opposite direction (Fig. 4). All mesenchymal tissues surrounding SOX2-deficient AFG epithelial tubes developed as respiratory organ mesenchyme rather than esophageal mesenchyme. Thus, interactions between epithelial and mesenchymal components during AFG development appear to be bidirectional and dependent on the developmental stage. The underlying mechanisms through which epithelial regional identity determines the characters of the surrounding mesenchyme remain to be elucidated. In a previous study, SOX2-deficient AFG failed to express some Wnt antagonists (Trisno et al., 2018), whereas in other studies, the anteroposterior organization of the gut tube depended on retinoic acid signaling (Bayha et al., 2009; Wang et al., 2006). Hence, the regulation of these signals by epithelial tissues may contribute to the coordinated development of epithelial and mesenchymal components of the AFG.

## Experimental procedures

### Mouse and mouse embryos

Knock-in mice expressing CreERT2 with the specificity of *FoxA2* (*Foxa2*^*nEGFP-CreERT2*^) (Imuta et al., 2013) or *ROSA26* (Gt*(ROSA)26Sor*^*tm1(cre/ERT2)Alj*^) (Cheng et al., 2010) were crossed with mice carrying a floxed *Sox2* allele (Fig. S1) and bred for several generations in C57BL/6;DBA mixed background to achieve homozygosity at both loci. Five mg of tamoxifen in peanut oil was orally administered to pregnant females on 7.5 and 8.5 dpc, and embryos were collected at the indicated stages. Animal experiments were performed at Kyoto Sangyo University, Nagoya University, Osaka University, and Tokushima University in accordance with the animal experimentation regulations of the respective institutions.

### Histological analyses

Specimens were fixed in 4% paraformaldehyde and processed for histological analyses. Immunofluorescence staining was done using antibodies listed in Table S1. Whole-mount *in situ* hybridization of isolated gut tubes was conducted under the conditions indicated in Table S2, followed by cryosectioning. Paraffin sections of embryos fixed with Bouin’s fixative were stained with hematoxylin and eosin (H&E). Photoimages were taken using Axioplan 2 (Zeiss) or an FV3000 laser microscope (Olympus).

## Acknowledgments

We thank Hiroshi Sasaki and Alexandra Joyner for granting the use of their *Foxa2*^*nEGFP-CreERT2*^ and Gt*(ROSA)26Sor*^*tm1(cre/ERT2)Alj*^ mice, respectively. This study was supported by Grants-in-Aid for Scientific Research JP19K16184 to MT and JP26251024 and JP17H03680 to HK.

